# Molecular epidemiology of carbapenemase-producing *Acinetobacter* spp. from Israel, 2001-2006: earliest report of *bla*_NDM_ predating the oldest known *bla*_NDM_-positive strains

**DOI:** 10.1101/2022.06.03.494778

**Authors:** Frédéric Grenier, Vincent Baby, Sarah Allard, Félix Heynemand, Simon Lévesque, Richard Sullivan, Hannah L. Landecker, Paul G. Higgins, Sébastien Rodrigue, Louis-Patrick Haraoui

## Abstract

**Background:** Carbapenem-resistant *Acinetobacter baumannii* (CRAb) is a WHO priority 1 critical pathogen. Despite early emergence of elevated CRAb rates in Israel, limited molecular data from this location are available. We searched for carbapenemases among 198 clinical *Acinetobacter* spp. from Israel between 2001 and 2006.

**Methods:** Strains from 3 archives underwent whole-genome sequencing (Illumina NovaSeq on all, MinION on a subset) and computational analyses: assembly (Unicycler), annotation (prokka), identification (Kraken, *rpoB* similarity), search for carbapenemases (ResFinder, BLDB curation).

**Findings:** *A. baumannii* (Ab) represented 179 (90·4%) *Acinetobacter* spp. Eighty-four Ab (46·9%) carried a carbapenemase: 38 (45·2%) *bla*_OXA-72_ (*bla*_OXA-24-like_); 28 (33·3%) *bla*_OXA-23-like_ (20 *bla*_OXA-23_ and 8 *bla*_OXA-225_); 18 (21·5%) *bla*_OXA-58_ (16 from 2001-2). Carbapenemase rates increased yearly from 2002 (32%) to 2006 (67%). Eight species of non-*baumannii Acinetobacter* (NbA) accounted for 19 isolates (9·6%). Two of three *A. junii* contained *bla*_OXA-58_, one of which, Ajun-H1-3, isolated in January 2004, also possessed *bla*_NDM-1_. The pNDM-Ajun-H1-3 plasmid matched numerous NDM-positive plasmids reported from 2005 onwards in *Acinetobacter* spp. as well as *Enterobacterales*.

**Interpretation:** We assessed carbapenemase diversity among *Acinetobacter* spp. in Israel from 2001-2006. Findings in Ab predate observations elsewhere: rapidly rising carbapenemase rates, driven by *bla*_OXA-23-like_ and *bla*_OXA-24-like_ genes replacing *bla*_OXA-58_. Among NbA, an *A. junii* isolated in 2004 carried *bla*_NDM-1_, making it the earliest NDM-positive isolate reported to date, preceding those from 2005 in India. Further research into *bla*_NDM_’s emergence is warranted, in order to shed light on the evolution and spread of this and other antibiotic-resistance genes.

**Funding:** Centre de recherche Charles-Le Moyne; Department of Microbiology and Infectious Diseases, Faculty of Medicine and Health Sciences, Université de Sherbrooke; Fonds de recherche du Québec – Santé; New Frontiers in Research Fund Grant NFRFE-2019-00444; CIFAR-Azrieli Global Scholars Program.

## Introduction

Antimicrobial resistance (AMR) is a major global public health issue. AMR deaths in 2019 surpassed HIV and malaria in terms of infectious disease-related mortality, being associated with 4·95 million deaths, including 1·27 million directly attributable to bacterial AMR^1^. In 2018, the World Health Organization (WHO) issued its priority list for discovery, research, and development of new treatments for antibiotic-resistant bacteria. All priority 1 critical pathogens – *Acinetobacter baumannii* (Ab), *Pseudomonas aeruginosa* and *Enterobacteriaceae* – share a common trait: resistance to carbapenems, the most recent class in the penicillin family of antibiotics^2^. Carbapenem-resistant Ab (CRAb) ranks 4^th^ in AMR deaths worldwide among pathogen-drug combinations, behind methicillin-resistant *Staphylococcus aureus*, multidrug-resistant tuberculosis and 3^rd^ generation cephalosporin-resistant *E. coli*^1^.

Carbapenemases, enzymes that confer resistance to carbapenems and other related drug classes, account for the most common mechanism through which bacteria become resistant to these antibiotics. One of these, New Delhi metallo-beta-lactamase (NDM), encoded by the *bla*_NDM_ gene, has become the most prevalent carbapenem-resistance gene among clinical isolates worldwide despite being reported more recently than the other predominant carbapenemases^3^. While *bla*_NDM_ likely originated in an “*Acinetobacter* background”^4,5^, CRAb seldom carries this gene, more commonly relying on acquired oxacillinases (*bla*_OXA_ genes)^6^. Among these, *bla*_OXA-23_ and related enzymes have become predominant, surpassing other common OXA groups such as *bla*_OXA-24_ and *bla*_OXA-58_.

One of the global regions initially affected by the rise and broad spread of carbapenem resistance in *Acinetobacter* spp. was the Middle East. Reports of a rapidly expanding incidence of carbapenem-resistance among *Acinetobacter* clinical isolates date back to the 1990’s in Israel^7^. Early during Operation Iraqi Freedom, the U.S. army noted alarming rates of CRAb infections among troops injured in combat^8^, largely due to the dissemination of *bla*_OXA-23_-carrying Ab (Patrick McGann, personal communication). The earliest NDM-positive *Acinetobacter* strain reported from the Middle East is an *A. pittii* found in Turkey in 2006^9^, one year after the oldest known isolates from India^4,5^.

A recent review of the molecular mechanisms of CRAb analysed 3575 publicly-available genomes of Ab up until April 2019^6^. Surprisingly, not a single isolate was labelled as originating from Israel. Given the known early, elevated rates of CRAb in Israel, the apparent role of *Acinetobacter* spp. in the emergence of *bla*_NDM_, and the lack of formal retrospective molecular analyses done among isolates from the early 2000’s, we searched for *bla*_NDM_ and other carbapenemases among 198 *Acinetobacter* clinical isolates from Israel between 2001 and 2006.

In this paper, we report the earliest NDM-positive isolate reported to date: a *bla*_NDM-1_-carrying *Acinetobacter junii* isolated in 2004 from a blood culture in a hematology patient hospitalized at Chaim Sheba Medical Center in Tel Hashomer, Israel. We highlight important trends seen across this collection, both in terms of clonal lineages, and the molecular mechanisms responsible for carbapenem-resistance in both Ab and non-*baumannii Acinetobacter* (NbA) in Israel. Understanding the origins and spread of antibiotic resistance, including the highly disseminated *bla*_NDM_ gene, sheds light on the processes promoting the emergence and dissemination of antibiotic-resistance genes, and impacts policy aimed at mitigating the drivers of AMR.

## Methods

### Bacterial isolates

We obtained 198 clinical *Acinetobacter* spp. isolated in Israel between 2001 and 2006 from three archives: 1) directly from Chaim Sheba Medical Center, Tel Hashomer, Israel (n= 140) referred to as hospital 1 (H1); 2) from JMI Laboratories who, as part of their SENTRY Antimicrobial Surveillance Program, still held 37 isolates also from Chaim Sheba Medical Center, distinct from those we obtained directly from the medical center (also accounted as H1); and 21 isolates from International Health Management Associates inc. (IHMA). These 21 isolates were collected from two different anonymized hospitals (H2, n=7; and H3, n=14). Strains are identified according to their genus and species name (A for *Acinetobacter* and the first three letters of their species), followed by the archive source (labelled H1, H2 or H3), ending with a number referring to the chronological order of isolation for each species/archive combination, with the number 1 referring to the earliest isolates. Finally, we also obtained from the Barcelona Institute for Global Health Foundation (ISGlobal) two NDM-positive NbA isolates for comparative purposes, an *A. pittii* (JVAP02) and an *A. lactucae* (JVAP01), both isolated in Turkey in 2006 and 2009 respectively^9,10^.

### Whole Genome Sequencing

All isolates underwent short-read sequencing. DNA libraries were prepared from extracted gDNA using the NEBNext Ultra II FS DNA Library Prep Kit for Illumina (NEB). DNA was purified and size selected using Ampure XP beads (Beckman Coulter) and quantified using Quant-it PicoGreen dsDNA assay (Thermo Fisher). The quality and size distribution of the DNA was assessed on a Fragment Analyzer using the HS NGS Fragment Kit (Agilent). The pooled samples were then sequenced on a NovaSeq6000 (Illumina) as PE250 by the McGill Genome Center (https://www.mcgillgenomecentre.ca/).

A subset of strains also underwent long-read DNA sequencing. Extracted gDNA was treated with the NEBNext Ultra II End Repair/dA-Tailing Module (NEB). Then barcodes from the Native Barcoding Expansion 1-12 & 13-24 from Oxford Nanopore Technologies (ONT) were ligated using the NEBNext Ultra II Ligation Module (NEB). DNA was purified using Ampure XP beads (Beckman Coulter). The DNA from different barcoded samples was pooled and the adapter AMII (ONT) was ligated using the NEBNext Ultra II Ligation Module (NEB). DNA was purified using Ampure XP beads (Beckman Coulter), followed by sequencing with a R10.4 MinION Flow Cell using a MinIon Mk1B (ONT).

The sequencing reads are deposited in genbank under BioProject number XXXXXXXXX.

### Data analysis/bioinformatics

For Illumina reads, the quality assessment and trimming was done using fastp 0·21·0 with --cut_right --cut_window_size 4 --cut_mean_quality 20 --length_required 30 --detect_adapter_for_pe^11^. Samples with less than 20X coverage were resequenced. Assemblies were made using Unicycler 0·4·9^12^ using the trimmed Illumina short reads and ONT long reads when available. Contigs were filtered to retain only those above 500 bp. Taxonomic identification was made on the assemblies using Kraken 2 (2·0·9-beta)^13^. Initially unresolved species identification (reported as *Acinetobacter sp*. by Kraken 2) were adjudicated using *rpoB* sequence similarity, and by JspeciesWS^14^ when further confirmation was considered necessary. Antibiotic resistance genes were found using ResFinder 3·0^15^. Detected *bla*_OXA_ variants were curated using BLDB^16^. Presence or absence of *bla*_OXA-51-like_ genes further contributed to discriminating between Ab and NbA.

Two distinct multi-locus sequence typing (MLST) schemes exist for Ab, known as Oxford and Institut Pasteur (STo and STp)^17,18^, with the latter being used in this paper. The ST of each strain was determined using mlst 2·11 (Seemann T, mlst Github https://github.com/tseemann/mlst) which made use of the PubMLST website (https://pubmlst.org/)^19^. *A. baumannii* core genome MLST (cgMLST)^20^ was obtained using chewBBACA 2·8·5^21^ and rendered with GrapeTree 2.2 using Standard Neighbour Joining^22^. Assemblies were annotated with Prokka 1·14·5^23^ using the additional databases Pfam, TIGRFAM, and the *bla*_OXA_ variants present in BLDB^16^. Plasmid alignments were made using AliTV^24^.

## Results

### Overview

A collection of 198 *Acinetobacter* spp. isolates from Israeli hospitals was assembled from three archives (see Table 1 in Supplementary Material for a list of isolates, their main characteristics, as well as sequencing technology used and outputs obtained). *A. baumannii* (Ab) accounted for 179 of 198 (90·4%) of the strains. The number of isolates varied between years, from a minimum of n=16 from 2001, to a maximum of n=62 in 2004, with an average of close to 30 isolates/year. Given that the vast majority of isolates are Ab, data are segregated between Ab and Non-*baumannii Acinetobacter* (NbA). Their distribution according to species identification, year of isolation, *bla*_OXA_ content, and Pasteur MLST (STp) is shown in Figures 1 and 2.

**Figure 1.**
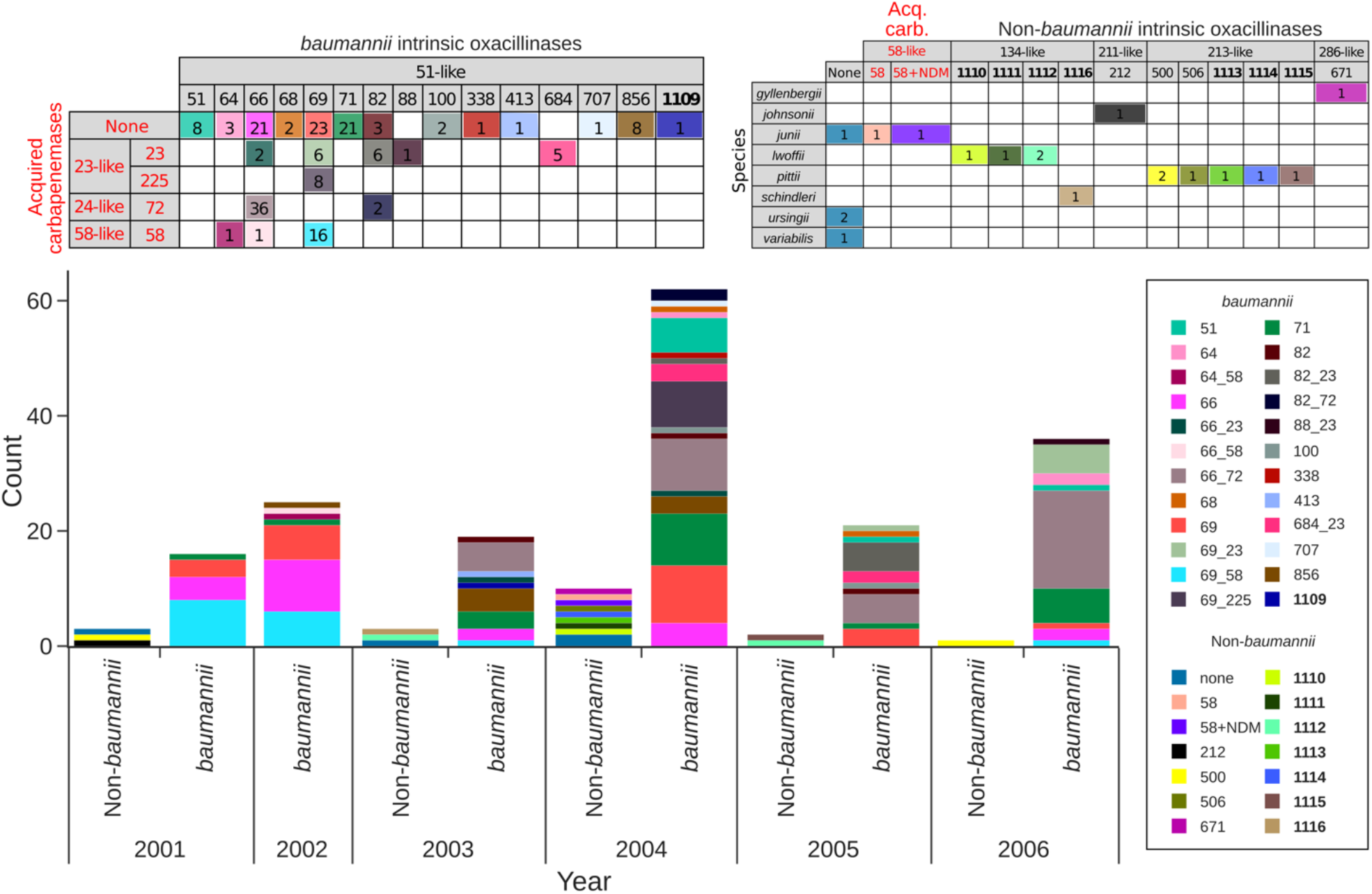
Distribution of *bla*_OXA_ genes among *Acinetobacter* spp. from Israel between 2001 and 2006. Data are segregated between *A. baumannii* and Non-*baumannii Acinetobacter*. The two tables present combinations of intrinsic oxacillinases (indicated in black), with, when present, acquired carbapenemases (in red). Novel oxacillinases initially described as part of this study (***bla***_**OXA-1109-1116**_) are in bold. Yearly counts highlight shifting trends in *bla*_OXA_ genes.

**Figure 2.**
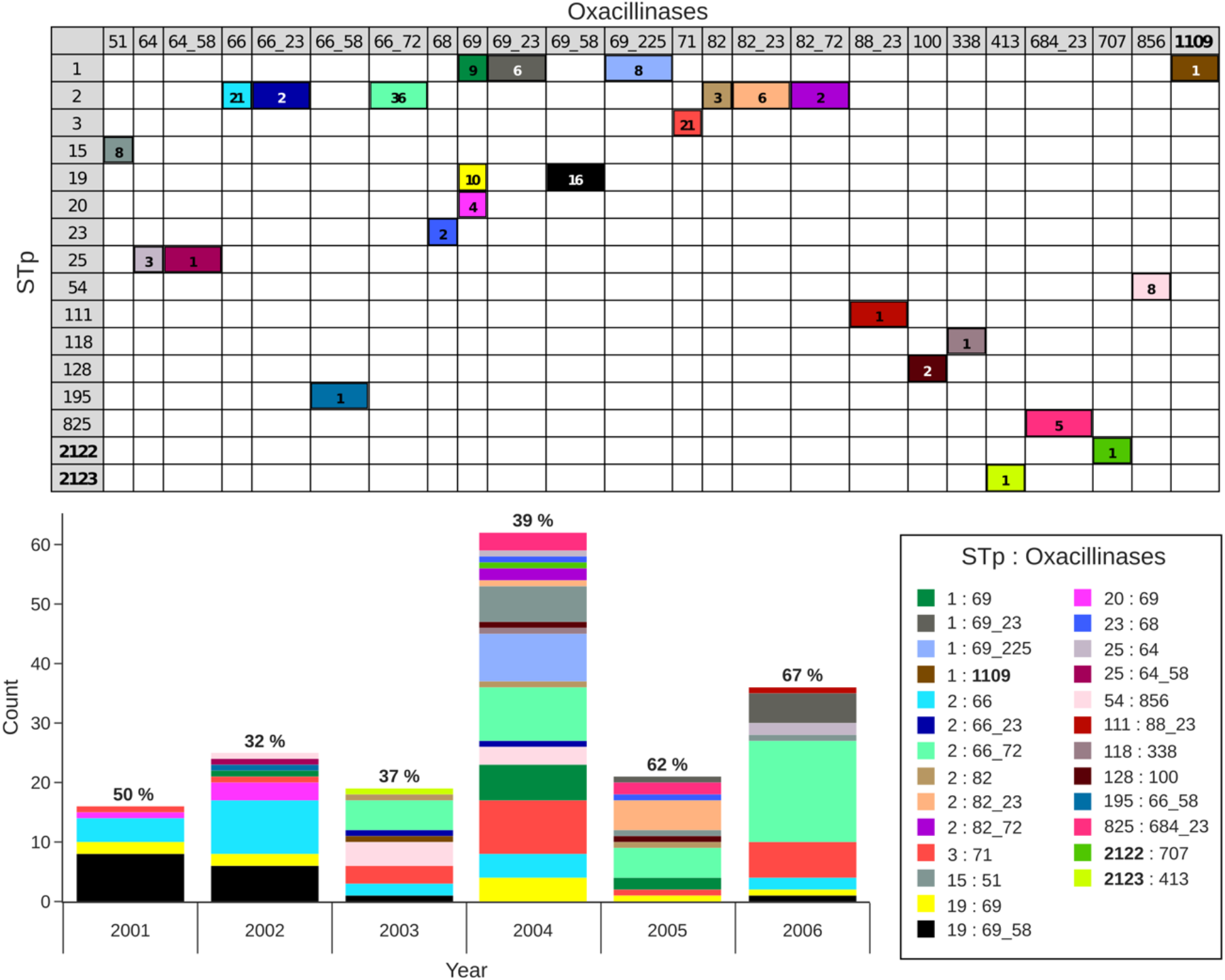
Pasteur MLST (STp) distribution among *Acinetobacter baumannii* according to *bla*_OXA_ gene content, and by study year. Yearly percentage of isolates with acquired carbapenemases are indicated above each annual count. Novel STp (**2122 and 2123**) and oxacillinase (***bla***_**OXA-1109**_) initially described as part of this study are in bold.

### *Acinetobacter baumannii (Ab)* collection

#### *bla*_OXA_ analysis

Distribution of *bla*_OXA_ genes is presented in Figure 1. All Ab are known to carry a *bla*_OXA-51-like_ gene that does not usually confer resistance to carbapenems unless overexpressed. The four most common *bla*_OXA-51-like_ genes representing 81% of Ab isolates (145/179) were, in decreasing order: i) *bla*_OXA-66_: 60 (33·5% overall, 59/60 of which are of STp2, the other being STp195), including 39 (65%) combined with another *bla*_OXA_ (36/39 with the *bla*_OXA-24-like_ variant *bla*_OXA-72_,); ii) *bla*_OXA-69_: 53 (29·6%), 31 (58·5%) combined with an additional *bla*_OXA_ (16/31 with *bla*_OXA-58_); iii) *bla*_OXA-71_ (21, 11·7%, all belonging to STp3), none of which carried an additional *bla*_OXA_ gene; and iv) *bla*_OXA-82_ (11, 6·1%, all belonging to STp2), 8 found alongside an acquired oxacillinase. Ten out of eleven *bla*_OXA-82_ were identified in the latter half of the study (2004-6), along with a single isolate from 2003 (none from 2001 and 2002). Eleven other *bla*_OXA-51-like_ genes were found, all in low numbers (range 1 to 8), including a novel gene variant, *bla*_OXA-1109_. In addition to the links between clones and *bla*_OXA_ genes noted above (STp2 and *bla*_OXA-66_; STp3 and *bla*_OXA-71_), certain unique associations between *bla*_OXA-51-like_ genes and sequence types were noted: i) 8 isolates with *bla*_OXA-856_ all belonging STp54; ii) 5 isolates with *bla*_OXA-684_ all linked to STp825 (all of which also carried *bla*_OXA-23_).

Overall, 95 (53·1%) Ab carried no additional oxacillinase beyond their *bla*_OXA-51-like_ gene. None carried more than two *bla*_OXA_ genes. Among the 84 isolates (46·9%) with a second *bla*_OXA_ gene, we identified four genes from all three oxacillinase subfamilies most commonly associated with carbapenem resistance in Ab: i) 38 (45·2%) *bla*_OXA-72_ (*bla*_OXA-24-like_), none from 2001-2; ii) 28 (33·3%) belonging to the *bla*_OXA-23-like_ subfamily (20 *bla*_OXA-23_, 71·4%, and 8 *bla*_OXA-225_, 28·6%, the latter all found within STp1 isolates); iii) 18 (21·5%) *bla*_OXA-58_, the bulk of which (16/18, 88·9%) were identified during the first two years of the study (2001-2), giving way to the two other subfamilies during the following years. The 8 *bla*_OXA-225_ were all identified in 2004, possibly as part of an outbreak, as they were not seen prior to or in the years that followed. The percentage of Ab isolates with acquired *bla*_OXA_ carbapenemases increased every year from 2002 (32%) to 2006 (67%), effectively more than doubling over a five-year period (Figure 2).

#### Pasteur Institute MLST Scheme (STp)

Ab STp distribution according to *bla*_OXA_ gene content, and by study year, is presented in Figure 2. The four most common sequence types, STp1, 2, 3 and 19 accounted for 78·8% of Ab isolates (141/179), close to half of which (70, 39·1%) are STp2, belonging to international clone 2 (IC2), the predominant clone worldwide. Over half of these (38, or 54·3%) carry *bla*_OXA-72_, a *bla*_OXA-24-like_ enzyme; of the remaining 32 STp2, 8 (11·4%) contain *bla*_OXA-23_, and 24 carry no other oxacillinase besides their *bla*_OXA-51-like_ gene. Other common Pasteur sequence types, presented in decreasing order, were: STp19 (26, 14·5%), all carrying *bla*_OXA-69_ as their *bla*_OXA-51-like_ gene, with 16 (61·5%) of these also possessing *bla*_OXA-58_; STp1 (24, 13·4%), 23 of these also containing *bla*_OXA-69_ as their *bla*_OXA-51-like_ gene (14, 58·3% also carrying a *bla*_OXA-23-like_ gene), and one attributed to the newly designated *bla*_OXA-1109_ (Abau-H1-57); STp3 (21, 11·7%) all possessing *bla*_OXA-71_. Twelve other STp profiles accounted for 38 further Ab isolates, all in single digits (range 1-8), including two new STp types, 2122 (Abau-H1-72) and 2123 (Abau-H1-59). The majority of STp19 were identified in the first two years of the study (2001-2), coinciding with the observation noted above regarding its association with *bla*_OXA-58_ also predominant early in the study. STp2 was the predominant sequence type every year except 2001.

#### cgMLST assessment

Neighbour joining trees derived from the cgMLST scheme^20^ are shown in Figures 3 and 4. cgMLST results correlate well with the STp distribution (Figure 3A). Intrinsic oxacillinases tend to cluster with certain STp (Figure 3B), as described above. The most common STp were distributed throughout the study period, except STp19 which predominated in 2001-2.

**Figure 3.**
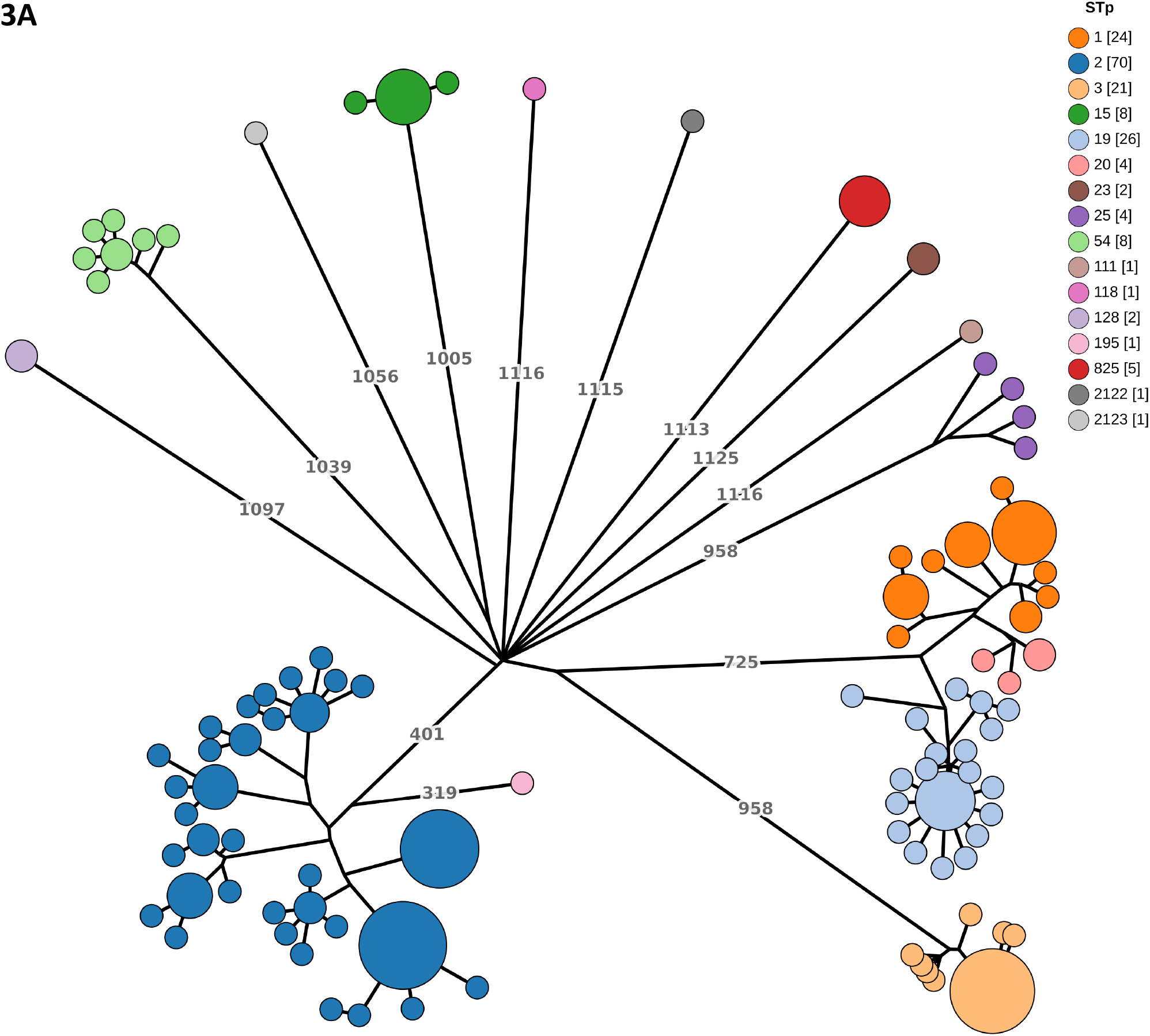
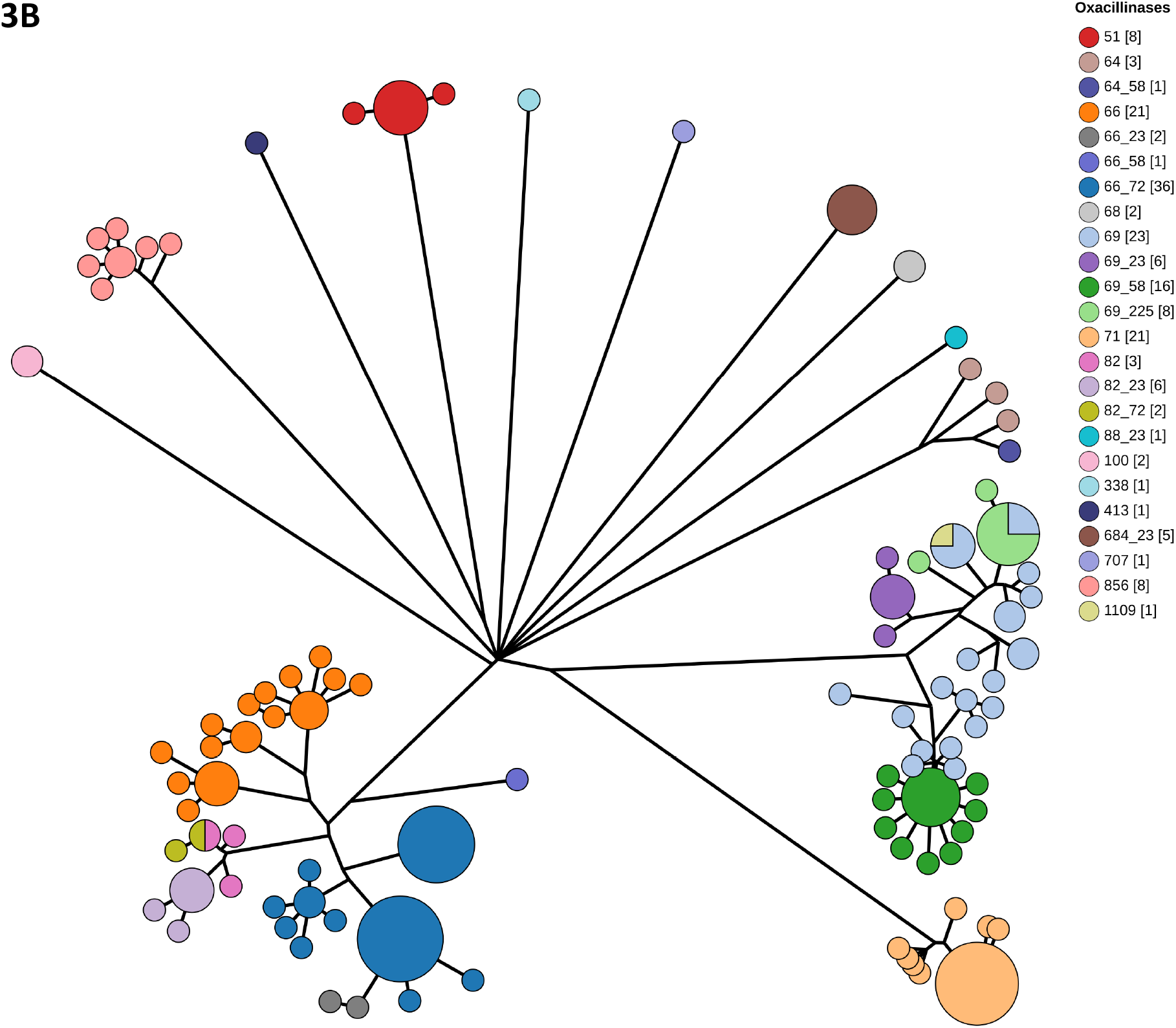
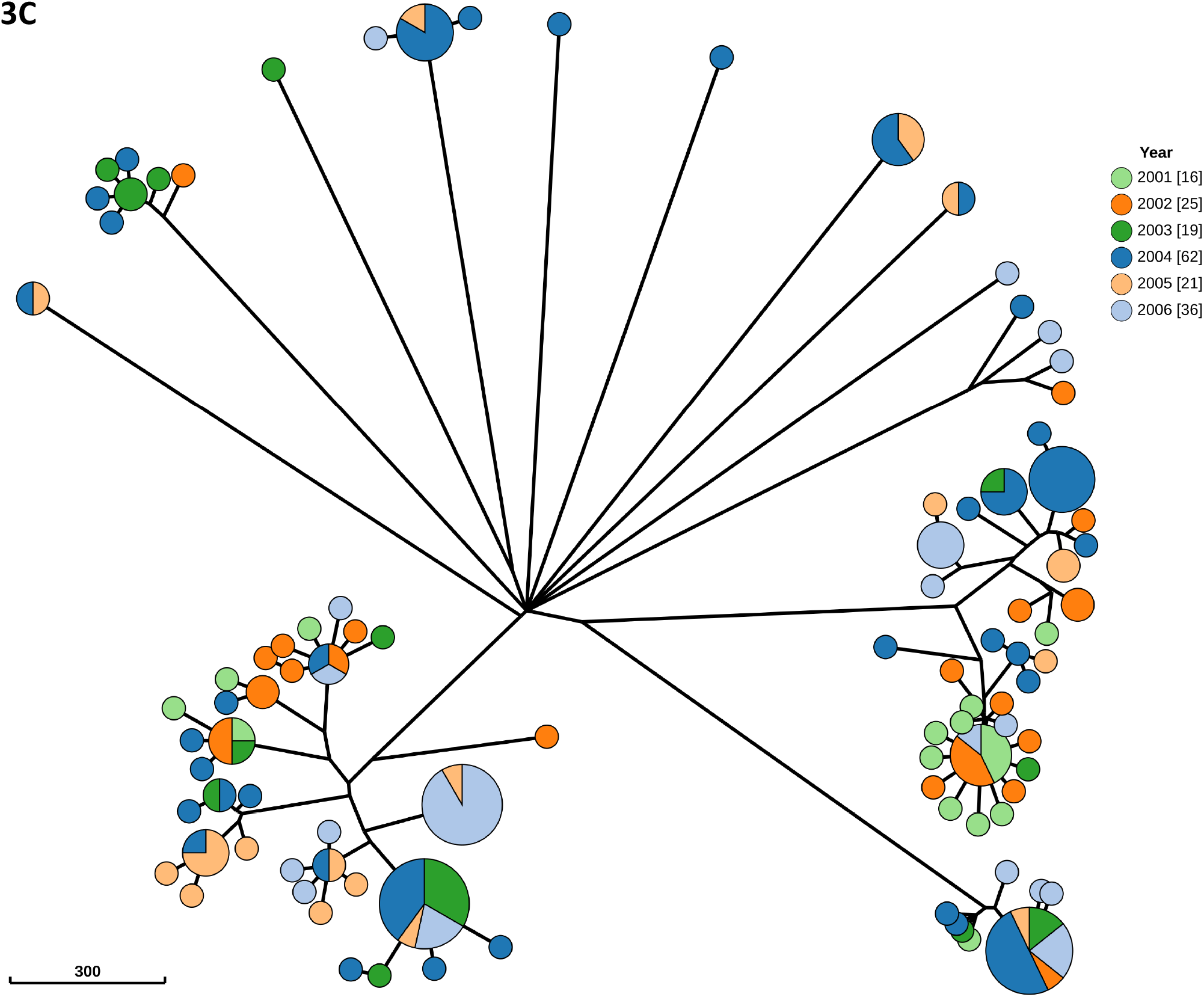
Neighbour joining trees (linear scale) derived from cgMLST of 179 *A. baumannii*. Strains or clusters of highly similar ones are colored according to **(A)** Pasteur MLST (STp), **(B)** *bla*_OXA_ gene content, and **(C)** year of isolation. Number of strains according to each variable are presented in brackets. Node distances of 300 alleles or more are indicated in **(A)**.

**Figure 4.**
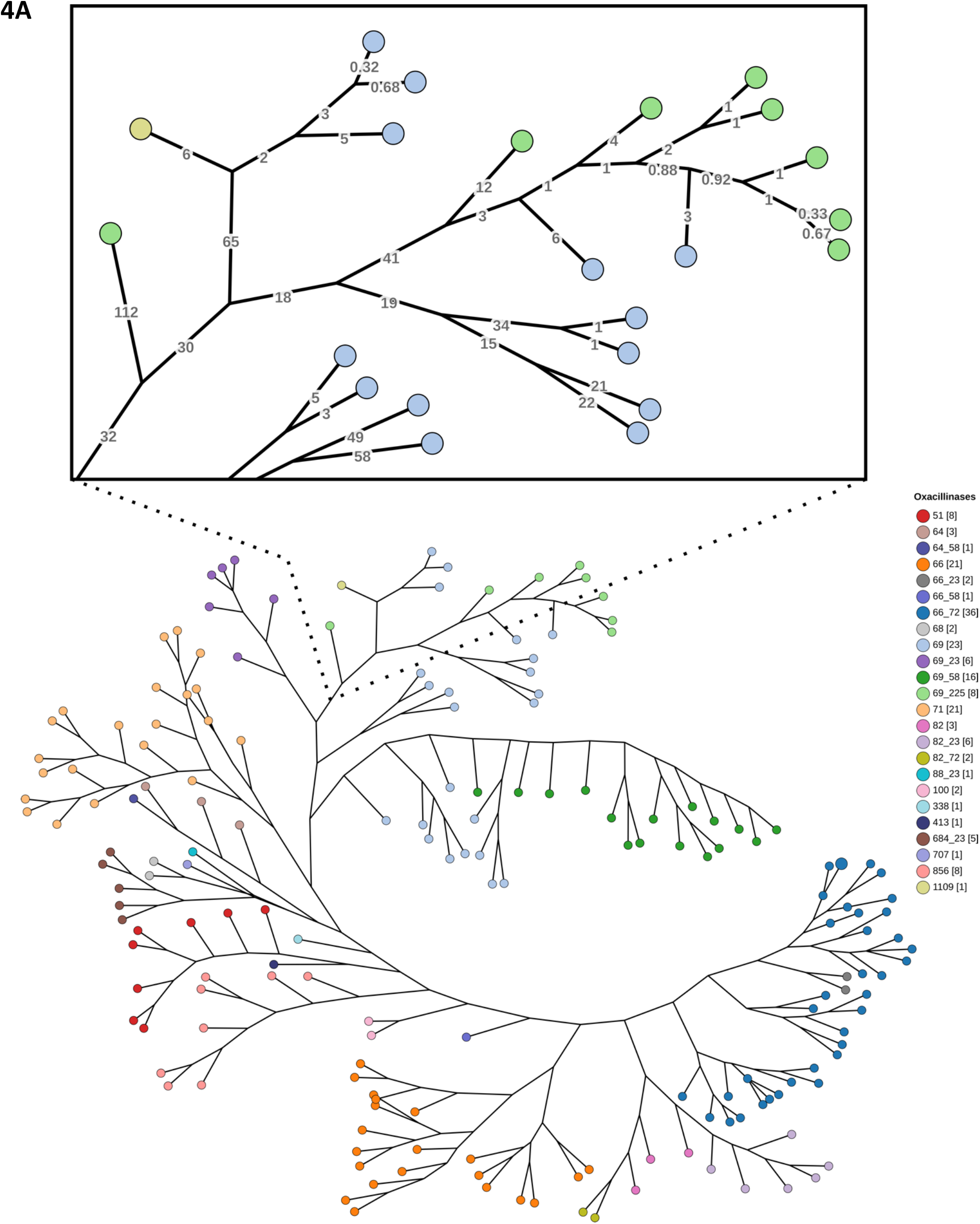
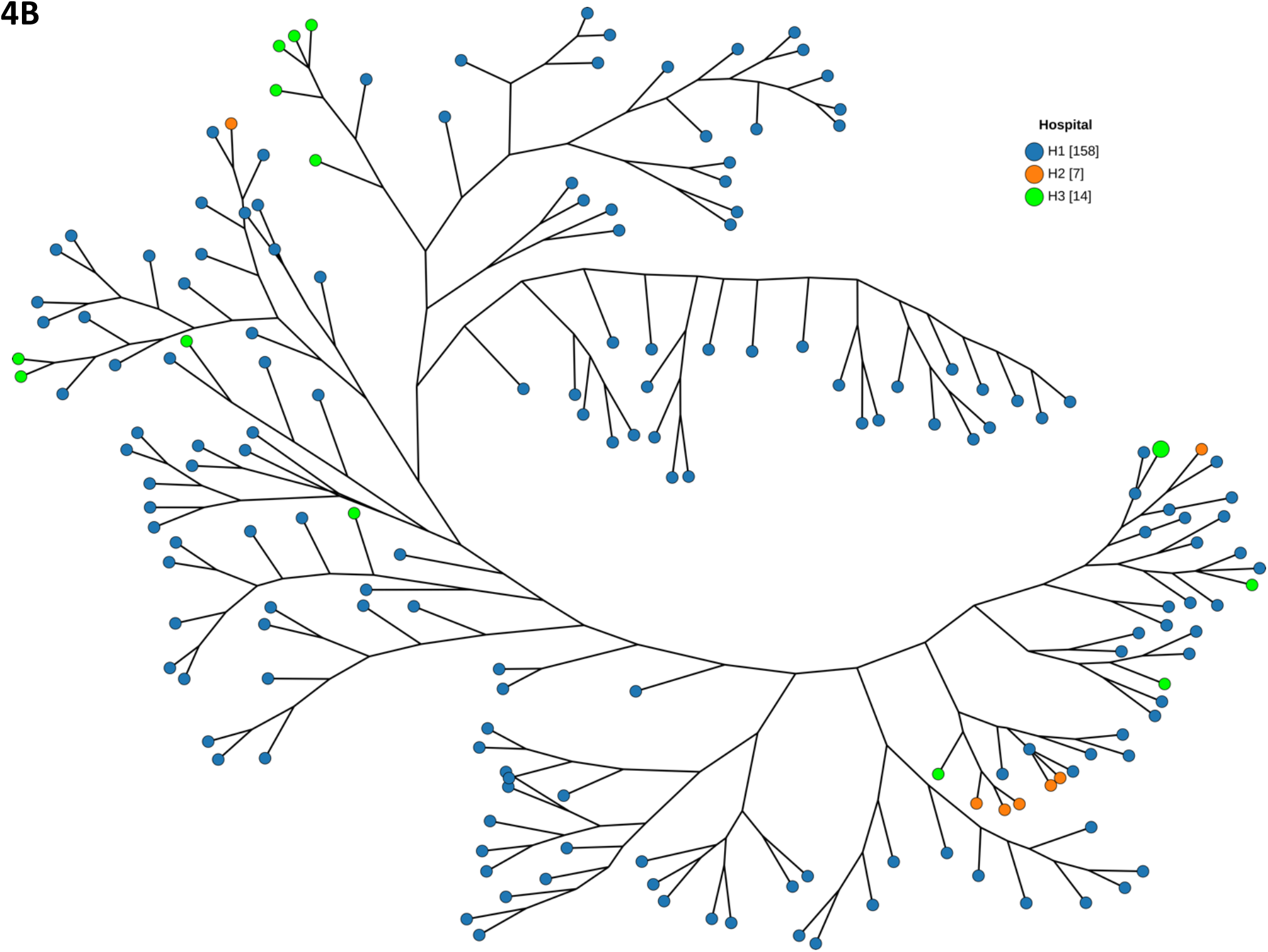
Neighbour joining trees (log scale) derived from cgMLST of 179 *A. baumannii*. Strains, or clusters of highly similar ones, are colored according to **(A)** *bla*_OXA_ gene content, and **(B)** source hospital. The close-up in panel **(A)** presents genetic differences among eight STp1 isolates carrying *bla*_OXA-69_ and *bla*_OXA-225_ from 2004 (light green) along with comparisons to other STp1 strains lacking acquired carbapenemases, including Abau-H1-57 (yellow-green) carrying the newly described *bla*_OXA-51-like_ gene *bla*_OXA-1109_. Number of strains according to each variable are presented in brackets.

In the case of the eight STp1 isolates carrying *bla*_OXA-69_ and *bla*_OXA-225_ found exclusively in 2004, Figure 4A lends support to the hypothesis that most of these isolates were part of an outbreak. They also happen to cluster with other H1-derived STp1 isolates with *bla*_OXA-69_ that do not carry the *bla*_OXA-225_ acquired carbapenemase (all from 2004), and with a single isolate from 2003 (Abau-H1-57) that carries the newly described *bla*_OXA-51-like_ gene *bla*_OXA-1109_. No other outbreak could be definitively ascertained except for possibly a cluster of 5 STp825 isolates from H1 spanning 2004 and 2005 that all carried *bla*_OXA-684_ as their *bla*_OXA-51-like_ gene, as well as *bla*_OXA-23_ (Figure 3). They differed significantly from the other isolates analyzed. Both STp2122 and 2123 reported for the first time in this study (Abau-H1-72 and Abau-H1-59 respectively), were unrelated and also differed significantly from all the other isolates. Some isolates from the anonymized H2 and H3 hospitals contained very few allelic differences to others from H1, Chaim Sheba Medical Center, (Figure 4B). These genetically-similar isolates from different hospitals were found among the dominant international clones (STp1 and STp2) that tend to circulate globally. Therefore we could not definitively rule in or out that H2 or H3 could also represent Chaim Sheba Medical Center.

#### Non-*baumannii Acinetobacter* (NbA) description

Nineteen isolates (9·6%) belonging to eight species were NbA, all from H1 (Figure 1). The predominant species were *A. pittii* (n=6), *A. lwoffii* (n=4), *A. junii* (n=3), *A. ursingii* (n=2), and four other species each with a single representative: *A. gyllenbergii, A. johnsonnii, A. schindleri*, and *A. variabilis*. Thirteen of 19 isolates carried an intrinsic oxacillinase, among which 7 novel *bla*_OXA_ genes were identified in 8 isolates: 4 *A. lwoffii, bla*_OXA-1110_ (Alwo-H1-2), *bla*_OXA-1111_ (Alwo-H1-3), and *bla*_OXA-1112_ (Alwo-H1-1 and Alwo-H1-4), all *bla*_OXA-134-like_; 3 *A. pittii, bla*_OXA-1113_ (Apit-H1-2), *bla*_OXA-1114_ (Apit-H1-3), and *bla*_OXA-1115_ (Apit-H1-5), all *bla*_OXA-213-like_; one *A. schindleri, bla*_OXA-1116_ (Asch-H1-1), also *bla*_OXA-134-like_. No *bla*_OXA_ genes were found in the two *A. ursingii* and the *A. variabilis*.

Among three unrelated *A. junii*, a species not known to carry an intrinsic oxacillinase, two contained a single copy of *bla*_OXA-58_ on two different plasmids. Strain Ajun-H1-3, isolated in a blood culture from a hematology patient in January 2004, also possessed *bla*_NDM-1_ on plasmid pNDM-Ajun-H1-3 which was distinct from the same isolate’s *bla*_OXA-58_-bearing plasmid. Ajun-H1-3 is the earliest known *bla*_NDM_ isolate to date. It is the sole NDM-positive isolate of the dataset. No records were accessible to obtain further details about the patient from whom it was recovered.

In pNDM-Ajun-H1-3, *bla*_NDM_ was found on a Tn*125* transposon, flanked by two full copies of IS*Aba*125, as has been often reported among the earliest NDM-positive isolates. A third copy of IS*Aba*125 was found on the same 49kb plasmid (exact size: 49,163 base pairs). This plasmid was found to match numerous other NDM-positive plasmids termed pNDM-BJ01-like, based on their similarity to a plasmid initially reported in an *A. lwoffii* strain from November 2010 in China^25^. Since this initial description, pNDM-BJ01-like plasmids have been shown to be widely circulating among *Acinetobacter* spp., particularly among NbA, including: pNDM-40-1 in an *A. bereziniae* from India from 2005^4,5^; pNDM-JVAP02 (*A. pittii* from Turkey from 2006)^26^; pNDM-JVAP01 (*A. lactucae* also from Turkey from 2009)^27^; numerous *Acinetobacter* spp. from China from 2009 onwards^28–30^, as well as elsewhere around the world. pNDM-Ajun-H1-3 is 99·99% similar over 97% of pNDM-40-1’s sequence, and 99·98% similar over 97% of pNDM-BJ01’s sequence. pNDM-BJ01-like plasmids have also been identified in Enterobacterales clinical isolates in China and in Mexico^31,32^. Finally, they have been shown to be transferable through conjugation experiments from *Acinetobacter* spp. to *E. coli*^31^. pNDM-BJ01-like plasmid alignments including pNDM-Ajun-H1-3, pNDM-JVAP01 and pNDM-JVAP02 are presented in Figure 5.

**Figure 5.**
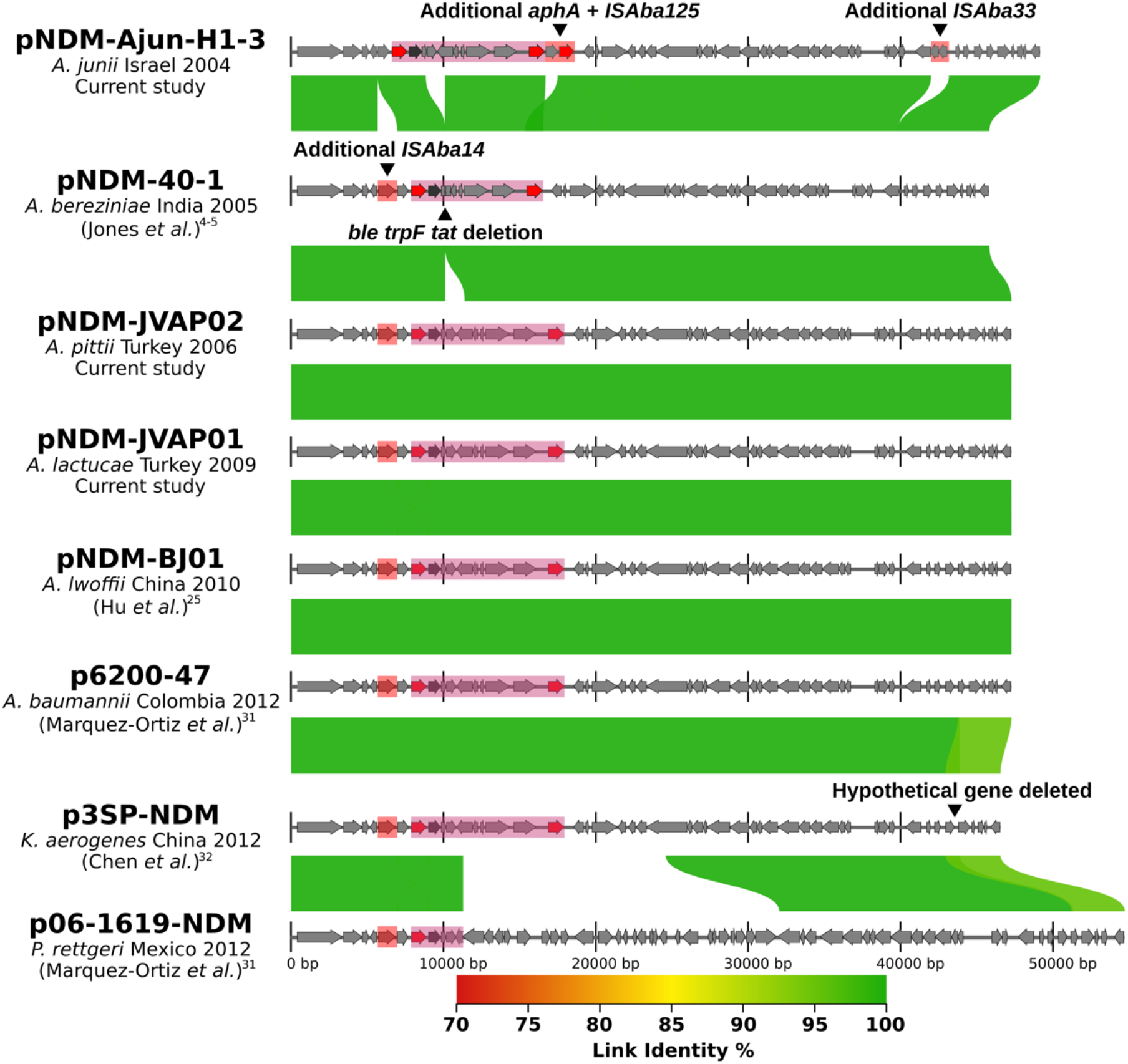
Alignment of certain BJ01-like NDM-positive plasmids in *Acinetobacter* spp. and Enterobacterales. Plasmids are presented in chronological order from top to bottom, starting with the earliest known description of *bla*_NDM_, reported as part of this study. Alignments demonstrate dissemination of highly conserved plasmids. The species identification, country and year of isolation are listed underneath the plasmid name. Below these are the references from which the plasmid sequences were accessed. Although data for pNDM-JVAP01 were already available^27^, the assembly presented here is derived from our own sequencing.

## Discussion

Carbapenem-resistant Ab poses a significant and growing risk to human health worldwide. According to a recent and large systematic analysis using 2019 data, CRAb ranks fourth on the list of deadliest AMR pathogen-drug threats^1^.

We retrospectively assessed the diversity of carbapenemases and clonal patterns among *Acinetobacter* spp. in Israel from the early 21^st^ century, a critical period in terms of the emergence, shifting predominance and spread of carbapenemases, notably oxacillinases and *bla*_NDM_. While Israeli institutions were among the earliest to signal a rise in carbapenem-resistance among *Acinetobacter* spp., starting in the 1990’s^7^, little to no molecular epidemiological data has been available from that country.

Analysis of 179 Ab isolates collected between 2001 and 2006 shows several significant trends in sequence types and genetic distribution of carbapenemases, particularly *bla*_OXA_ genes. Notably, these results are contemporaneous and in some cases predate similar phenomena observed in other countries many years later: i) a predominance of certain sequence types, the majority belonging to international clone 2 (STp2), over half of which carry carbapenemases; ii) rapidly rising rates of carbapenem resistance, most notably driven by the dissemination of *bla*_OXA-23-like_ and *bla*_OXA-24-like_ genes, gradually outpacing clones carrying *bla*_OXA-58_^6^. The rate of carbapenemase-positive Ab more than doubled between 2002 and 2006, from 32% to 67%. The rate for the first year of the study (2001) was 50%, an elevated and discordant result with the overall trend observed in the years that followed. This is possibly a result of biased archiving of a smaller number of isolates (n=16), with calculations being susceptible to greater shifts in the relative abundance of genes.

Among 179 Ab strains, we identified two new Ab STp profiles, 2122 and 2123. Along with the Ab strains, we analysed 19 NbA (close to 10% of the archival collection), belonging to eight species. Among the 198 total *Acinetobacter* spp. isolates, we identified eight new intrinsic *bla*_OXA_ genes, *bla*_OXA-1109_ (*bla*_OXA-51-like_) within one Ab, and 7 *bla*_OXA_ genes (*bla*_OXA-1110 to 1116_) among 8 NbA (4 *A. lwoffii*, 3 *A. pittii* and one *A. schindleri*).

Among NbA, isolate Ajun-H1-3 isolated from a blood culture in January 2004 was the only isolate of the entire collection to carry two acquired carbapenemases, *bla*_OXA-58_ and *bla*_NDM-1_. To our knowledge, Ajun-H1-3 is the earliest NDM-positive isolate reported, predating the seven NDM-positive *Acinetobacter* spp. (six *A. baumannii* and one *A. bereziniae*) from Tamil Nadu from 2005^4,5^. An *A. junii* also carrying *bla*_OXA-58_ and *bla*_NDM-1_ was isolated from surveillance swabs from a five-day old Palestinian girl in October 2012, more than eight years later^33^. The latter isolate is no longer available to carry out whole genome sequencing to compare it to Ajun-H1-3 (Musa Y. Hindiyeh, personal communication).

*bla*_NDM-1_ was initially reported in a 2008 poster at ICAAC^34^, followed by a 2009 publication describing the initial two isolates known to carry this gene, a *Klebsiella pneumoniae* and an *Escherichia coli*^35^. These were identified in 2008 in a single patient who was receiving care in Sweden following a trip to India during which he had been hospitalized in Ludhiana and in New Delhi. Since this initial description, *bla*_NDM_ has become the most prevalent carbapenemase gene worldwide, finding its way into numerous γ-Proteobacteria including Enterobacterales, *Acinetobacter* spp. and *Pseudomonas* spp.^3^. Over thirty variants of this enzyme exist, with new ones still being described (https://card.mcmaster.ca/, last accessed May 26, 2022). *bla*_NDM_ has been implicated in human infections, identified in domestic and farm animals, and spread to environments devoid of human presence such as High Arctic soil ecosystems^36–38^. Retrospective analyses have consistently shown the earliest NDM-positive strains in a given geographical region to be *Acinetobacter* spp. Until now, the earliest NDM-positive strains were identified in 2005 from India^4,5^. A study on 22 Gram-negative strains from India from 2000 did not find any evidence of this gene^39^.

The significance of the NDM-positive-Ajun-H1-3 identified in this study lies in its implications for our understanding of the origins of *bla*_NDM_. Molecular analyses point to the origins of *bla*_NDM_ within an *Acinetobacter* background^4,5^. Retrospective work on bacterial archives supports this claim, with growing evidence pointing to NbA as the likely source of *bla*_NDM_, as does this study. The fact that the gene was found in *A. junii* further supports the hypothesis that *bla*_NDM_ originated in bacteria with a predominantly environmental footprint, namely NbA, a phenomenon that has been observed with numerous other antibiotic-resistance genes of significant concern to human health (e.g. *bla*_CTX-M_, *bla*_OXA-23_, *bla*_OXA-48_)^40–42^.

Two critical and contemporary shifts occurring over the last decades can help explain the pivotal role of *Acinetobacter* spp. in the *bla*_NDM_ story: i) a shifting chemical milieu, both in the clinic and in the environment^43^, followed by the introduction of carbapenem antibiotics in the 1980’s; and ii) the intrinsic capacity of many *Acinetobacter* spp. to exhibit resistance to carbapenems, allowing them to gain an increasing foothold in human healthcare settings. The *Acinetobacter* mobilome, already capable of circulating mobile genetic elements (MGEs) among *Acinetobacter* spp., increasingly began to overlap with the mobilome of more common human pathogens, namely *Enterobacterales*^44^.

A recent study assessing the role of MGEs in the global dissemination of *bla*_NDM_ estimated that this carbapenemase emerged on a Tn*125* transposon, the same genetic construct we report being present in pNDM-Ajun-H1-3, prior to “1990, and possibly well back into the mid-twentieth century”^45^. If this latter scenario were to be confirmed, this would indicate that *bla*_NDM_ emerged prior to the commercialization of carbapenem antibiotics. This is not unlike the identification of the acquired carbapenemase *bla*_OXA-58_ in an *Acinetobacter indicus* isolated in 1953, decades before carbapenems were used^46^.

In conclusion, the identification of a *bla*_NDM_-positive strain from Israel, predating what were previously the earliest known isolates from India in 2005, supports the need for further work in investigating the origins of this widespread and problematic gene. Authors of a recent study maintain that the Indian subcontinent is still the likely geographical region where *bla*_NDM_ emerged despite their modelling pointing to its origins occurring decades prior to its first earliest known presence in India in 2005^45^. Our findings caution against premature conclusions as to *bla*_NDM_’s origins, understanding of which is still incomplete. The socio-environmental conditions leading to *bla*_NDM_’s emergence may have been quite different than those that contributed to its dissemination, as has been demonstrated in many other instances^47–49^. *bla*_NDM_ could very well have originated elsewhere than in India, before disseminating widely within its borders, in neighbouring countries and globally. Focusing on NbA and environmental archives from the 20^th^ and early 21^st^ centuries may yield additional clues, helping to understand the factors that led to its emergence and prevent similar issues from arising in the future.

## Acknowledgments

we would like to acknowledge and thank Chaim Sheba Medical Center (Drs Gill Smollan and Sharon Amit), JMI Laboratories (Helio Sader) and IHMA (Meredith Hackel) as the sources from which the Israeli isolates were obtained. We would also like to thank ISGlobal (Prof. Ignasi Roca) for sharing two NDM-positive strains isolated in Turkey (JVAP01 and JVAP02).

## The authors have no conflicts of interest to declare

Louis-Patrick Haraoui is funded by the Centre de recherche Charles-Le Moyne; the Department of Microbiology and Infectious Diseases, Faculty of Medicine and Health Sciences, Université de Sherbrooke; the Fonds de recherche du Québec – Santé; a New Frontiers in Research Fund Grant, NFRFE-2019-00444; and by the CIFAR-Azrieli Global Scholars Program.

## References

1 Murray CJ, Ikuta KS, Sharara F, et al. Global burden of bacterial antimicrobial resistance in 2019: a systematic analysis. The Lancet 2022; 399: 629–55.

2 Tacconelli E, Carrara E, Savoldi A, et al. Discovery, research, and development of new antibiotics: the WHO priority list of antibiotic-resistant bacteria and tuberculosis. The Lancet Infectious Diseases 2018; 18: 318–27.

3 Logan LK, Weinstein RA. The Epidemiology of Carbapenem-Resistant Enterobacteriaceae: The Impact and Evolution of a Global Menace. The Journal of Infectious Diseases 2017; 215: S28–36.

4 Jones LS, Toleman MA, Weeks JL, Howe RA, Walsh TR, Kumarasamy KK. Plasmid Carriage of bla_NDM-1_ in Clinical Acinetobacter baumannii Isolates from India. Antimicrob Agents Chemother 2014; 58: 4211–3.

5 Jones LS, Carvalho MJ, Toleman MA, et al. Characterization of Plasmids in Extensively Drug-Resistant Acinetobacter Strains Isolated in India and Pakistan. Antimicrob Agents Chemother 2015; 59: 923–9.

6 Hamidian M, Nigro SJ. Emergence, molecular mechanisms and global spread of carbapenem-resistant Acinetobacter baumannii. Microbial Genomics 2019; 5. DOI:10.1099/mgen.0.000306.

7 Simhon A, Rahav G, Shazberg G, Block C, Bercovier H, Shapiro M. Acinetobacter baumannii at a Tertiary-Care Teaching Hospital in Jerusalem, Israel. J Clin Microbiol 2001; 39: 389–91.

8 Centers for Disease Control and Prevention (CDC). Acinetobacter baumannii infections among patients at military medical facilities treating injured U.S. service members, 2002-2004. MMWR Morb Mortal Wkly Rep 2004; 53: 1063–6.

9 Roca I, Mosqueda N, Altun B, Espinal P, Akova M, Vila J. Molecular characterization of NDM-1-producing Acinetobacter pittii isolated from Turkey in 2006. Journal of Antimicrobial Chemotherapy 2014; 69: 3437–8.

10 Espinal P, Mosqueda N, Telli M, et al. Identification of NDM-1 in a Putatively Novel Acinetobacter Species (“NB14”) Closely Related to Acinetobacter pittii. Antimicrob Agents Chemother 2015; 59: 6657–60.

11 Chen S, Zhou Y, Chen Y, Gu J. fastp: an ultra-fast all-in-one FASTQ preprocessor. Bioinformatics 2018; 34: i884–90.

12 Wick RR, Judd LM, Gorrie CL, Holt KE. Unicycler: Resolving bacterial genome assemblies from short and long sequencing reads. PLoS Comput Biol 2017; 13: e1005595.

13 Wood DE, Lu J, Langmead B. Improved metagenomic analysis with Kraken 2. Genome Biol 2019; 20: 257.

14 Richter M, Rosselló-Móra R, Oliver Glöckner F, Peplies J. JSpeciesWS: a web server for prokaryotic species circumscription based on pairwise genome comparison. Bioinformatics 2016; 32: 929–31.

15 Zankari E, Hasman H, Cosentino S, et al. Identification of acquired antimicrobial resistance genes. Journal of Antimicrobial Chemotherapy 2012; 67: 2640–4.

16 Naas T, Oueslati S, Bonnin RA, et al. Beta-lactamase database (BLDB) – structure and function. Journal of Enzyme Inhibition and Medicinal Chemistry 2017; 32: 917–9.

17 Bartual SG, Seifert H, Hippler C, Wisplinghoff H, Rodriguez-Valera F. Development of a Multilocus Sequence Typing Scheme for Characterization of Clinical Isolates of. J CLIN MICROBIOL 2005; 43: 9.

18 Diancourt L, Passet V, Nemec A, Dijkshoorn L, Brisse S. The Population Structure of Acinetobacter baumannii: Expanding Multiresistant Clones from an Ancestral Susceptible Genetic Pool. PLoS ONE 2010; 5: e10034.

19 Jolley KA, Bray JE, Maiden MCJ. Open-access bacterial population genomics: BIGSdb software, the PubMLST.org website and their applications. Wellcome Open Res 2018; 3: 124.

20 Higgins PG, Prior K, Harmsen D, Seifert H. Development and evaluation of a core genome multilocus typing scheme for whole-genome sequence-based typing of Acinetobacter baumannii. PLoS ONE 2017; 12: e0179228.

21 Silva M, Machado MP, Silva DN, et al. chewBBACA: A complete suite for gene-by-gene schema creation and strain identification. Microbial Genomics 2018; 4. DOI:10.1099/mgen.0.000166.

22 Zhou Z, Alikhan N-F, Sergeant MJ, et al. GrapeTree: visualization of core genomic relationships among 100,000 bacterial pathogens. Genome Res 2018; 28: 1395–404.

23 Seemann T. Prokka: rapid prokaryotic genome annotation. Bioinformatics 2014; 30: 2068–9.

24 Ankenbrand MJ, Hohlfeld S, Hackl T, Förster F. AliTV—interactive visualization of whole genome comparisons. PeerJ Computer Science 2017; 3: e116.

25 Hu H, Hu Y, Pan Y, et al. Novel Plasmid and Its Variant Harboring both a bla_NDM-1_ Gene and Type IV Secretion System in Clinical Isolates of Acinetobacter lwoffii. Antimicrob Agents Chemother 2012; 56: 1698–702.

26 Roca I, Mosqueda N, Altun B, Espinal P, Akova M, Vila J. Molecular characterization of NDM-1-producing Acinetobacter pittii isolated from Turkey in 2006. Journal of Antimicrobial Chemotherapy 2014; 69: 3437–8.

27 Cosgaya C, Marí-Almirall M, Van Assche A, et al. Acinetobacter dijkshoorniae sp. nov., a member of the Acinetobacter calcoaceticus–Acinetobacter baumannii complex mainly recovered from clinical samples in different countries. International Journal of Systematic and Evolutionary Microbiology 2016; 66: 4105–11.

28 Fu Y, Du X, Ji J, Chen Y, Jiang Y, Yu Y. Epidemiological characteristics and genetic structure of blaNDM-1 in non-baumannii Acinetobacter spp. in China. Journal of Antimicrobial Chemotherapy 2012; 67: 2114–22.

29 Huang T-W, Lauderdale T-L, Liao T-L, et al. Effective transfer of a 47 kb NDM-1-positive plasmid among Acinetobacter species. J Antimicrob Chemother 2015; 70: 2734–8.

30 Zhang R, Hu Y-Y, Yang X-F, et al. Emergence of NDM-producing non-baumannii Acinetobacter spp. isolated from China. Eur J Clin Microbiol Infect Dis 2014; 33: 853–60.

31 Marquez-Ortiz RA, Haggerty L, Olarte N, et al. Genomic Epidemiology of NDM-1-Encoding Plasmids in Latin American Clinical Isolates Reveals Insights into the Evolution of Multidrug Resistance. Genome Biology and Evolution 2017; 9: 1725–41.

32 Chen Z, Li H, Feng J, et al. NDM-1 encoded by a pNDM-BJ01-like plasmid p3SP-NDM in clinical Enterobacter aerogenes. Front Microbiol 2015; 6. DOI:10.3389/fmicb.2015.00294.

33 Regeen H, Al-Sharafa-Kittaneh D, Kattan R, Al-Dawodi R, Marzouqa H, Hindiyeh MY. First Report of bla _NDM_ and bla_OXA-58_ Coexistence in Acinetobacter junii. J Clin Microbiol 2014; 52: 3492–3.

34 Yong D, Giske CG, Toleman M, Walsh TR. A novel subgroup metallo-beta-lactamase (MBL), NDM-1 emerges in Klebsiella pneumoniae (KPN) from India. In: 48th Annual ICAAC/IDSA 46th Annual Meeting, Washington DC. 2008: C1–105.

35 Yong D, Toleman MA, Giske CG, et al. Characterization of a New Metallo-β-Lactamase Gene, bla_NDM-1_, and a Novel Erythromycin Esterase Gene Carried on a Unique Genetic Structure in Klebsiella pneumoniae Sequence Type 14 from India. Antimicrob Agents Chemother 2009; 53: 5046–54.

36 Wu W, Feng Y, Tang G, Qiao F, McNally A, Zong Z. NDM Metallo-β-Lactamases and Their Bacterial Producers in Health Care Settings. Clin Microbiol Rev 2019; 32. DOI:10.1128/CMR.00115-18.

37 McCann CM, Christgen B, Roberts JA, et al. Understanding drivers of antibiotic resistance genes in High Arctic soil ecosystems. Environment International 2019; 125: 497–504.

38 Wang B, Sun D. Detection of NDM-1 carbapenemase-producing Acinetobacter calcoaceticus and Acinetobacter junii in environmental samples from livestock farms. Journal of Antimicrobial Chemotherapy 2015; 70: 611–3.

39 Castanheira M, Deshpande L, Woosley L, Prochaska R, Jones R. Retrospective Search for NDM-1 Reveals Possible Indian Origin of DIM-1 Metallo-beta-lactamase. ; : 1.

40 Pitout JDD, Peirano G, Kock MM, Strydom K-A, Matsumura Y. The Global Ascendency of OXA-48-Type Carbapenemases. Clin Microbiol Rev 2019; 33: e00102–19.

41 Poirel L, Figueiredo S, Cattoir V, Carattoli A, Nordmann P. Acinetobacter radioresistens as a Silent Source of Carbapenem Resistance for Acinetobacter spp. Antimicrob Agents Chemother 2008; 52: 1252–6.

42 Humeniuk C, Arlet G, Gautier V, Grimont P, Labia R, Philippon A. ^N^_L_-Lactamases of Kluyvera ascorbata, Probable Progenitors of Some Plasmid-Encoded CTX-M Types. ANTIMICROB AGENTS CHEMOTHER 2002; 46: 5.

43 Landecker H. Antimicrobials before antibiotics: war, peace, and disinfectants. Palgrave Commun 2019; 5: 45.

44 Haraoui L-P. Networked collective microbiomes and the rise of subcellular ‘units of life’. Trends in Microbiology 2022; 30: 112–9.

45 Acman M, Wang R, van Dorp L, et al. Role of mobile genetic elements in the global dissemination of the carbapenem resistance gene blaNDM. Nat Commun 2022; 13: 1131.

46 Bonnin RA, Dortet L, Naas T. Acquired carbapenemase in Acinetobacter during the pre-antibiotic era. The Lancet Microbe 2021; 2: e137.

47 Buchholz U, Bernard H, Werber D, et al. German Outbreak of Escherichia coli O104:H4 Associated with Sprouts. N Engl J Med 2011; 365: 1763–70.

48 Njamkepo E, Fawal N, Tran-Dien A, et al. Global phylogeography and evolutionary history of Shigella dysenteriae type 1. Nat Microbiol 2016; 1: 16027.

49 Tarequl Islam M, Alam M, Boucher Y. Emergence, ecology and dispersal of the pandemic generating Vibrio cholerae lineage. International Microbiology Official journal of the Spanish Society for Microbiology 2017; : 106–15.

