## Supplemental Table 1 for "Molecular epidemiology of carbapenemase-producing *Acinetobacter* spp. from Israel, 2001-2006: earliest report of *bla*_NDM_ predating the oldest known *bla*_NDM_-positive strains"

**Table 1. Main characteristics of 198 *Acinetobacter* spp. including sequencing technology used and outputs obtained.**
Isolate Ajun-H1-3 (p.7) is in red to indicate the presence of $bla_{NDM-1}$.

| Name | Year | STp | Species | Assembly type | Genome length | Contigs | Illumina coverage | ONT coverage | OXAs curated | OXA types curated |
|---|---|---|---|---|---|---|---|---|---|---|
| Abau-H1-1 | 2001 | 19 | *baumannii* | Illumina | 3802614 | 76 | 40,8 | | 58_69 | 51_58 |
| Abau-H1-2 | 2001 | 19 | *baumannii* | Illumina | 3843381 | 112 | 34,5 | | 58_69 | 51_58 |
| Abau-H1-3 | 2001 | 2 | *baumannii* | Illumina | 3980652 | 124 | 41,3 | | 66 | 51 |
| Abau-H1-4 | 2001 | 19 | *baumannii* | Illumina | 3880901 | 91 | 47,5 | | 69 | 51 |
| Abau-H1-5 | 2001 | 19 | *baumannii* | Illumina | 3847270 | 91 | 57,9 | | 58_69 | 51_58 |
| Abau-H1-6 | 2001 | 19 | *baumannii* | Illumina | 3848221 | 86 | 49,7 | | 58_69 | 51_58 |
| Abau-H1-7 | 2001 | 3 | *baumannii* | Illumina | 3880149 | 102 | 37,5 | | 71 | 51 |
| Abau-H1-8 | 2001 | 19 | *baumannii* | Illumina | 3809207 | 79 | 50,2 | | 58_69 | 51_58 |
| Abau-H1-9 | 2001 | 20 | *baumannii* | Illumina | 3985035 | 112 | 42,4 | | 69 | 51 |
| Abau-H1-10 | 2001 | 19 | *baumannii* | Illumina | 3847097 | 83 | 39,9 | | 58_69 | 51_58 |
| Abau-H1-11 | 2001 | 19 | *baumannii* | Illumina | 3844327 | 69 | 47,8 | | 69 | 51 |
| Abau-H1-12 | 2001 | 2 | *baumannii* | Illumina | 3995117 | 99 | 40,8 | | 66 | 51 |
| Abau-H1-13 | 2001 | 2 | *baumannii* | Illumina | 3969306 | 87 | 48,1 | | 66 | 51 |
| Abau-H1-14 | 2001 | 19 | *baumannii* | Illumina | 3858038 | 81 | 43,0 | | 58_69 | 51_58 |
| Abau-H1-15 | 2001 | 2 | *baumannii* | Illumina | 3983036 | 95 | 41,8 | | 66 | 51 |
| Abau-H1-16 | 2001 | 19 | *baumannii* | Illumina | 3788673 | 94 | 36,1 | | 58_69 | 51_58 |
| Abau-H1-17 | 2002 | 2 | *baumannii* | Illumina | 3964485 | 104 | 47,0 | | 66 | 51 |
| Abau-H1-18 | 2002 | 54 | *baumannii* | Illumina | 4148274 | 66 | 47,0 | | 856 | 51 |
| Abau-H1-19 | 2002 | 19 | *baumannii* | Illumina | 3810196 | 78 | 48,9 | | 58_69 | 51_58 |
| Abau-H1-20 | 2002 | 2 | *baumannii* | Illumina | 4041857 | 108 | 59,0 | | 66 | 51 |
| Abau-H1-21 | 2002 | 20 | *baumannii* | Illumina | 3913667 | 89 | 47,0 | | 69 | 51 |
| Abau-H1-22 | 2002 | 19 | *baumannii* | Illumina | 3832954 | 77 | 53,7 | | 58_69 | 51_58 |
| Abau-H1-23 | 2002 | 19 | *baumannii* | Illumina | 3843650 | 80 | 38,1 | | 58_69 | 51_58 |
| Abau-H1-24 | 2002 | 19 | *baumannii* | Illumina | 3881432 | 83 | 43,0 | | 58_69 | 51_58 |
| Abau-H1-25 | 2002 | 195 | *baumannii* | Hybrid | 4197988 | 5 | 47,4 | 108,3 | 58_66 | 51_58 |
| Abau-H1-26 | 2002 | 1 | *baumannii* | Illumina | 4038377 | 52 | 104,5 | | 69 | 51 |
| Abau-H1-27 | 2002 | 19 | *baumannii* | Illumina | 3695897 | 69 | 57,0 | | 69 | 51 |
| Abau-H1-28 | 2002 | 19 | *baumannii* | Illumina | 3852638 | 99 | 42,2 | | 58_69 | 51_58 |
| Abau-H1-29 | 2002 | 19 | *baumannii* | Illumina | 3883143 | 80 | 47,7 | | 58_69 | 51_58 |
| Abau-H1-30 | 2002 | 19 | *baumannii* | Illumina | 3915178 | 72 | 160,2 | | 69 | 51 |
| Abau-H1-31 | 2002 | 2 | *baumannii* | Illumina | 3925514 | 52 | 129,2 | | 66 | 51 |

| Name | Year | STp | Species | Assembly type | Genome length | Contigs | Illumina coverage | ONT coverage | OXAs curated | OXA types curated |
|---|---|---|---|---|---|---|---|---|---|---|
| Abau-H1-32 | 2002 | 2 | *baumannii* | Illumina | 4134641 | 134 | 54,7 | | 66 | 51 |
| Abau-H1-33 | 2002 | 2 | *baumannii* | Illumina | 3965886 | 104 | 68,1 | | 66 | 51 |
| Abau-H1-34 | 2002 | 20 | *baumannii* | Illumina | 4037408 | 72 | 47,6 | | 69 | 51 |
| Abau-H1-35 | 2002 | 2 | *baumannii* | Illumina | 4064566 | 88 | 43,8 | | 66 | 51 |
| Abau-H1-36 | 2002 | 2 | *baumannii* | Illumina | 3918570 | 93 | 42,8 | | 66 | 51 |
| Abau-H1-37 | 2002 | 3 | *baumannii* | Illumina | 3863928 | 48 | 49,7 | | 71 | 51 |
| Abau-H1-38 | 2002 | 20 | *baumannii* | Illumina | 4025477 | 75 | 50,5 | | 69 | 51 |
| Abau-H1-39 | 2002 | 2 | *baumannii* | Illumina | 3939230 | 111 | 62,5 | | 66 | 51 |
| Abau-H1-40 | 2002 | 25 | *baumannii* | Illumina | 4237426 | 52 | 58,3 | | 58_64 | 51_58 |
| Abau-H1-41 | 2002 | 2 | *baumannii* | Illumina | 4123482 | 119 | 49,0 | | 66 | 51 |
| Abau-H1-42 | 2003 | 19 | *baumannii* | Illumina | 3876249 | 103 | 30,9 | | 58_69 | 51_58 |
| Abau-H1-43 | 2003 | 2 | *baumannii* | Illumina | 3955783 | 92 | 52,3 | | 66_72 | 51_24 |
| Abau-H1-44 | 2003 | 3 | *baumannii* | Illumina | 3861202 | 42 | 49,7 | | 71 | 51 |
| Abau-H1-45 | 2003 | 2 | *baumannii* | Illumina | 3953528 | 95 | 53,8 | | 66_72 | 51_24 |
| Abau-H1-46 | 2003 | 2 | *baumannii* | Illumina | 3900136 | 68 | 66,7 | | 23_66 | 51_23 |
| Abau-H1-47 | 2003 | 2 | *baumannii* | Illumina | 3953983 | 85 | 52,7 | | 66_72 | 51_24 |
| Abau-H1-48 | 2003 | 3 | *baumannii* | Illumina | 3865533 | 43 | 49,5 | | 71 | 51 |
| Abau-H1-49 | 2003 | 2 | *baumannii* | Illumina | 3954317 | 87 | 69,6 | | 66_72 | 51_24 |
| Abau-H1-50 | 2003 | 54 | *baumannii* | Illumina | 4262661 | 88 | 44,6 | | 856 | 51 |
| Abau-H1-51 | 2003 | 54 | *baumannii* | Illumina | 4295356 | 63 | 37,1 | | 856 | 51 |
| Abau-H1-52 | 2003 | 2 | *baumannii* | Illumina | 3896769 | 73 | 51,1 | | 66 | 51 |
| Abau-H1-53 | 2003 | 54 | *baumannii* | Illumina | 4194803 | 106 | 33,7 | | 856 | 51 |
| Abau-H1-54 | 2003 | 2 | *baumannii* | Illumina | 3919008 | 89 | 42,9 | | 66_72 | 51_24 |
| Abau-H1-55 | 2003 | 2 | *baumannii* | Illumina | 3861629 | 121 | 58,7 | | 82 | 51 |
| Abau-H1-56 | 2003 | 3 | *baumannii* | Illumina | 3859367 | 39 | 41,4 | | 71 | 51 |
| Abau-H1-57 | 2003 | 1 | *baumannii* | Illumina | 4023120 | 51 | 44,7 | | 1109 | 51 |
| Abau-H1-58 | 2003 | 2 | *baumannii* | Illumina | 3993771 | 85 | 38,6 | | 66 | 51 |
| Abau-H1-59 | 2003 | 2123 | *baumannii* | Illumina | 4062251 | 198 | 43,8 | | 413 | 51 |
| Abau-H1-60 | 2003 | 54 | *baumannii* | Illumina | 4188354 | 55 | 54,8 | | 856 | 51 |
| Abau-H1-61 | 2004 | 54 | *baumannii* | Illumina | 4204062 | 80 | 40,4 | | 856 | 51 |

| Name | Year | STp | Species | Assembly type | Genome length | Contigs | Illumina coverage | ONT coverage | OXAs curated | OXA types curated |
|---|---|---|---|---|---|---|---|---|---|---|
| Abau-H1-62 | 2004 | 2 | *baumannii* | Illumina | 3911753 | 75 | 56,0 | | 66_72 | 51_24 |
| Abau-H1-63 | 2004 | 1 | *baumannii* | Illumina | 4031269 | 76 | 48,1 | | 225_69 | 51_23 |
| Abau-H1-64 | 2004 | 3 | *baumannii* | Illumina | 3856439 | 42 | 56,8 | | 71 | 51 |
| Abau-H1-65 | 2004 | 54 | *baumannii* | Illumina | 4221041 | 106 | 51,1 | | 856 | 51 |
| Abau-H1-66 | 2004 | 2 | *baumannii* | Illumina | 3882820 | 96 | 51,5 | | 72_82 | 51_24 |
| Abau-H1-67 | 2004 | 15 | *baumannii* | Illumina | 4035600 | 61 | 43,1 | | 51 | 51 |
| Abau-H1-68 | 2004 | 825 | *baumannii* | Illumina | 3864497 | 48 | 46,7 | | 23_684 | 51_23 |
| Abau-H1-69 | 2004 | 23 | *baumannii* | Illumina | 3923522 | 42 | 47,9 | | 68 | 51 |
| Abau-H1-70 | 2004 | 19 | *baumannii* | Illumina | 3987927 | 77 | 46,9 | | 69 | 51 |
| Abau-H1-71 | 2004 | 3 | *baumannii* | Illumina | 3829734 | 54 | 50,0 | | 71 | 51 |
| Abau-H1-72 | 2004 | 2122 | *baumannii* | Illumina | 3745930 | 33 | 55,9 | | 707 | 51 |
| Abau-H1-73 | 2004 | 2 | *baumannii* | Illumina | 4057970 | 113 | 31,5 | | 66_72 | 51_24 |
| Abau-H1-74 | 2004 | 118 | *baumannii* | Illumina | 3733350 | 31 | 42,1 | | 338 | 51 |
| Abau-H1-75 | 2004 | 3 | *baumannii* | Illumina | 3832110 | 48 | 39,9 | | 71 | 51 |
| Abau-H1-76 | 2004 | 128 | *baumannii* | Illumina | 4037645 | 132 | 62,1 | | 100 | 51 |
| Abau-H1-77 | 2004 | 2 | *baumannii* | Illumina | 3972211 | 83 | 55,9 | | 66 | 51 |
| Abau-H1-78 | 2004 | 1 | *baumannii* | Illumina | 4014549 | 45 | 50,7 | | 69 | 51 |
| Abau-H1-79 | 2004 | 1 | *baumannii* | Illumina | 3957006 | 68 | 49,4 | | 225_69 | 51_23 |
| Abau-H1-80 | 2004 | 3 | *baumannii* | Illumina | 3856139 | 46 | 51,4 | | 71 | 51 |
| Abau-H1-81 | 2004 | 15 | *baumannii* | Illumina | 4035222 | 67 | 41,2 | | 51 | 51 |
| Abau-H1-82 | 2004 | 19 | *baumannii* | Illumina | 3915136 | 73 | 47,6 | | 69 | 51 |
| Abau-H1-83 | 2004 | 1 | *baumannii* | Illumina | 4019638 | 56 | 85,4 | | 69 | 51 |
| Abau-H1-84 | 2004 | 2 | *baumannii* | Illumina | 4057828 | 108 | 57,7 | | 66_72 | 51_24 |
| Abau-H1-85 | 2004 | 2 | *baumannii* | Illumina | 3901622 | 70 | 65,4 | | 23_66 | 51_23 |
| Abau-H1-86 | 2004 | 15 | *baumannii* | Illumina | 4026585 | 76 | 40,7 | | 51 | 51 |
| Abau-H1-87 | 2004 | 825 | *baumannii* | Illumina | 3860624 | 64 | 66,0 | | 23_684 | 51_23 |
| Abau-H1-88 | 2004 | 19 | *baumannii* | Illumina | 3977088 | 82 | 41,9 | | 69 | 51 |
| Abau-H1-89 | 2004 | 25 | *baumannii* | Illumina | 4239774 | 86 | 40,8 | | 64 | 51 |
| Abau-H1-90 | 2004 | 1 | *baumannii* | Illumina | 4027241 | 61 | 52,7 | | 69 | 51 |

| Name | Year | STp | Species | Assembly type | Genome length | Contigs | Illumina coverage | ONT coverage | OXAs curated | OXA types curated |
|---|---|---|---|---|---|---|---|---|---|---|
| Abau-H1-91 | 2004 | 3 | *baumannii* | Illumina | 3824300 | 47 | 42,6 | | 71 | 51 |
| Abau-H1-92 | 2004 | 3 | *baumannii* | Illumina | 3812542 | 39 | 43,4 | | 71 | 51 |
| Abau-H1-93 | 2004 | 2 | *baumannii* | Illumina | 3839472 | 94 | 53,9 | | 82 | 51 |
| Abau-H1-94 | 2004 | 2 | *baumannii* | Illumina | 3982167 | 58 | 120,2 | | 66 | 51 |
| Abau-H1-95 | 2004 | 3 | *baumannii* | Illumina | 3861529 | 36 | 82,2 | | 71 | 51 |
| Abau-H1-96 | 2004 | 2 | *baumannii* | Illumina | 3885227 | 93 | 54,9 | | 72_82 | 51_24 |
| Abau-H1-97 | 2004 | 1 | *baumannii* | Illumina | 4012943 | 48 | 49,5 | | 69 | 51 |
| Abau-H1-98 | 2004 | 15 | *baumannii* | Illumina | 3998496 | 67 | 50,4 | | 51 | 51 |
| Abau-H1-99 | 2004 | 1 | *baumannii* | Illumina | 4036189 | 68 | 67,0 | | 225_69 | 51_23 |
| Abau-H1-100 | 2004 | 2 | *baumannii* | Illumina | 3818108 | 82 | 72,8 | | 23_82 | 51_23 |
| Abau-H1-101 | 2004 | 1 | *baumannii* | Illumina | 3963021 | 56 | 52,8 | | 225_69 | 51_23 |
| Abau-H1-102 | 2004 | 15 | *baumannii* | Illumina | 3980334 | 58 | 36,3 | | 51 | 51 |
| Abau-H1-103 | 2004 | 825 | *baumannii* | Illumina | 3865716 | 49 | 55,5 | | 23_684 | 51_23 |
| Abau-H1-104 | 2004 | 3 | *baumannii* | Illumina | 3861840 | 48 | 44,7 | | 71 | 51 |
| Abau-H1-105 | 2004 | 2 | *baumannii* | Illumina | 3933663 | 65 | 67,2 | | 66_72 | 51_24 |
| Abau-H1-106 | 2004 | 1 | *baumannii* | Illumina | 4002388 | 56 | 60,6 | | 225_69 | 51_23 |
| Abau-H1-107 | 2004 | 2 | *baumannii* | Illumina | 4000536 | 105 | 61,7 | | 66_72 | 51_24 |
| Abau-H1-108 | 2004 | 1 | *baumannii* | Illumina | 4033979 | 64 | 60,1 | | 225_69 | 51_23 |
| Abau-H1-109 | 2004 | 3 | *baumannii* | Illumina | 3852696 | 50 | 42,2 | | 71 | 51 |
| Abau-H1-110 | 2004 | 19 | *baumannii* | Illumina | 3840214 | 86 | 47,5 | | 69 | 51 |
| Abau-H1-111 | 2004 | 1 | *baumannii* | Illumina | 3935315 | 181 | 30,7 | | 225_69 | 51_23 |
| Abau-H1-112 | 2004 | 1 | *baumannii* | Illumina | 3995306 | 58 | 46,5 | | 69 | 51 |
| Abau-H1-113 | 2004 | 2 | *baumannii* | Illumina | 4015844 | 115 | 46,7 | | 66_72 | 51_24 |
| Abau-H1-114 | 2004 | 2 | *baumannii* | Illumina | 4017658 | 107 | 47,6 | | 66_72 | 51_24 |
| Abau-H1-115 | 2004 | 2 | *baumannii* | Illumina | 3939344 | 126 | 41,9 | | 66 | 51 |
| Abau-H1-116 | 2004 | 1 | *baumannii* | Illumina | 4029527 | 71 | 49,3 | | 225_69 | 51_23 |
| Abau-H1-117 | 2004 | 54 | *baumannii* | Illumina | 4226046 | 91 | 41,4 | | 856 | 51 |
| Abau-H1-118 | 2004 | 2 | *baumannii* | Illumina | 4167261 | 129 | 39,1 | | 66 | 51 |
| Abau-H1-119 | 2004 | 1 | *baumannii* | Illumina | 3989156 | 45 | 40,3 | | 69 | 51 |

| Name | Year | STp | Species | Assembly type | Genome length | Contigs | Illumina coverage | ONT coverage | OXAs curated | OXA types curated |
|---|---|---|---|---|---|---|---|---|---|---|
| Abau-H1-120 | 2004 | 2 | *baumannii* | Illumina | 3918853 | 84 | 44,1 | | 66_72 | 51_24 |
| Abau-H1-121 | 2004 | 15 | *baumannii* | Illumina | 4021007 | 74 | 44,2 | | 51 | 51 |
| Abau-H1-122 | 2004 | 2 | *baumannii* | Illumina | 3844206 | 84 | 51,2 | | 66_72 | 51_24 |
| Abau-H1-123 | 2005 | 128 | *baumannii* | Illumina | 4030274 | 138 | 44,4 | | 100 | 51 |
| Abau-H1-124 | 2005 | 2 | *baumannii* | Illumina | 3879164 | 114 | 37,2 | | 23_82 | 51_23 |
| Abau-H1-125 | 2005 | 2 | *baumannii* | Illumina | 3872148 | 113 | 54,7 | | 23_82 | 51_23 |
| Abau-H1-126 | 2005 | 2 | *baumannii* | Illumina | 3881811 | 103 | 51,3 | | 23_82 | 51_23 |
| Abau-H1-127 | 2005 | 1 | *baumannii* | Illumina | 3987856 | 55 | 55,8 | | 69 | 51 |
| Abau-H1-128 | 2005 | 1 | *baumannii* | Illumina | 3987819 | 56 | 43,6 | | 69 | 51 |
| Abau-H1-129 | 2005 | 825 | *baumannii* | Illumina | 3856216 | 50 | 43,3 | | 23_684 | 51_23 |
| Abau-H1-130 | 2005 | 825 | *baumannii* | Illumina | 3863220 | 69 | 37,0 | | 23_684 | 51_23 |
| Abau-H1-131 | 2005 | 3 | *baumannii* | Illumina | 3821985 | 45 | 42,3 | | 71 | 51 |
| Abau-H1-132 | 2005 | 2 | *baumannii* | Illumina | 3882260 | 102 | 28,5 | | 23_82 | 51_23 |
| Abau-H1-133 | 2005 | 2 | *baumannii* | Illumina | 3957107 | 126 | 53,4 | | 66_72 | 51_24 |
| Abau-H1-134 | 2005 | 2 | *baumannii* | Illumina | 4003827 | 114 | 53,2 | | 66_72 | 51_24 |
| Abau-H1-135 | 2005 | 2 | *baumannii* | Illumina | 3884633 | 99 | 111,5 | | 82 | 51 |
| Abau-H1-136 | 2005 | 1 | *baumannii* | Illumina | 3999735 | 68 | 55,3 | | 23_69 | 51_23 |
| Abau-H1-137 | 2005 | 19 | *baumannii* | Illumina | 3985145 | 75 | 75,7 | | 69 | 51 |
| Abau-H1-138 | 2005 | 2 | *baumannii* | Illumina | 3880920 | 103 | 78,5 | | 23_82 | 51_23 |
| Abau-H1-139 | 2005 | 15 | *baumannii* | Illumina | 4033130 | 64 | 54,0 | | 51 | 51 |
| Abau-H1-140 | 2005 | 23 | *baumannii* | Illumina | 3928044 | 53 | 42,0 | | 68 | 51 |
| Abau-H1-141 | 2005 | 2 | *baumannii* | Illumina | 3949628 | 153 | 33,7 | | 66_72 | 51_24 |
| Abau-H1-142 | 2006 | 2 | *baumannii* | Illumina | 3909377 | 145 | 41,1 | | 66_72 | 51_24 |
| Abau-H1-143 | 2006 | 2 | *baumannii* | Illumina | 3910637 | 140 | 40,8 | | 66_72 | 51_24 |
| Abau-H1-144 | 2006 | 2 | *baumannii* | Illumina | 4014089 | 112 | 96,7 | | 66_72 | 51_24 |
| Abau-H1-145 | 2006 | 2 | *baumannii* | Illumina | 3918428 | 113 | 114,6 | | 66_72 | 51_24 |
| Abau-H1-146 | 2006 | 2 | *baumannii* | Illumina | 3813413 | 130 | 43,8 | | 66_72 | 51_24 |
| Abau-H1-147 | 2006 | 3 | *baumannii* | Illumina | 3820498 | 63 | 43,2 | | 71 | 51 |
| Abau-H1-148 | 2006 | 25 | *baumannii* | Illumina | 4083900 | 76 | 37,7 | | 64 | 51 |

| Name | Year | STp | Species | Assembly type | Genome length | Contigs | Illumina coverage | ONT coverage | OXAs curated | OXA types curated |
|---|---|---|---|---|---|---|---|---|---|---|
| Abau-H1-149 | 2006 | 2 | *baumannii* | Illumina | 4076717 | 110 | 39,5 | | 66_72 | 51_24 |
| Abau-H1-150 | 2006 | 2 | *baumannii* | Illumina | 3914872 | 115 | 38,5 | | 66_72 | 51_24 |
| Abau-H1-151 | 2006 | 2 | *baumannii* | Illumina | 3989572 | 65 | 104,9 | | 66 | 51 |
| Abau-H1-152 | 2006 | 19 | *baumannii* | Illumina | 3922332 | 82 | 44,7 | | 69 | 51 |
| Abau-H1-153 | 2006 | 3 | *baumannii* | Illumina | 3813720 | 63 | 32,6 | | 71 | 51 |
| Abau-H1-154 | 2006 | 3 | *baumannii* | Illumina | 3754011 | 91 | 36,6 | | 71 | 51 |
| Abau-H1-155 | 2006 | 19 | *baumannii* | Illumina | 3903477 | 78 | 66,5 | | 58_69 | 51_58 |
| Abau-H1-156 | 2006 | 111 | *baumannii* | Illumina | 4005485 | 91 | 31,4 | | 23_88 | 51_23 |
| Abau-H1-157 | 2006 | 2 | *baumannii* | Illumina | 3966204 | 112 | 36,6 | | 66_72 | 51_24 |
| Abau-H1-158 | 2006 | 2 | *baumannii* | Illumina | 3949015 | 112 | 54,5 | | 66 | 51 |
| Abau-H2-1 | 2005 | 2 | *baumannii* | Illumina | 3977702 | 100 | 45,3 | | 66_72 | 51_24 |
| Abau-H2-2 | 2005 | 2 | *baumannii* | Illumina | 3956048 | 95 | 43,0 | | 66_72 | 51_24 |
| Abau-H2-3 | 2006 | 2 | *baumannii* | Illumina | 3981880 | 81 | 114,4 | | 66_72 | 51_24 |
| Abau-H2-4 | 2006 | 2 | *baumannii* | Illumina | 3975748 | 99 | 39,5 | | 66_72 | 51_24 |
| Abau-H2-5 | 2006 | 2 | *baumannii* | Illumina | 4017042 | 103 | 141,2 | | 66_72 | 51_24 |
| Abau-H2-6 | 2006 | 2 | *baumannii* | Illumina | 3975367 | 96 | 43,1 | | 66_72 | 51_24 |
| Abau-H2-7 | 2006 | 3 | *baumannii* | Illumina | 3839292 | 35 | 71,7 | | 71 | 51 |
| Abau-H3-1 | 2006 | 1 | *baumannii* | Illumina | 3992048 | 71 | 111,4 | | 23_69 | 51_23 |
| Abau-H3-2 | 2006 | 15 | *baumannii* | Illumina | 3971322 | 84 | 21,0 | | 51 | 51 |
| Abau-H3-3 | 2006 | 2 | *baumannii* | Illumina | 3990012 | 102 | 31,6 | | 66_72 | 51_24 |
| Abau-H3-4 | 2006 | 2 | *baumannii* | Illumina | 3958734 | 78 | 50,0 | | 66_72 | 51_24 |
| Abau-H3-5 | 2006 | 2 | *baumannii* | Illumina | 3839803 | 86 | 27,7 | | 66_72 | 51_24 |
| Abau-H3-6 | 2006 | 1 | *baumannii* | Illumina | 3990757 | 67 | 50,8 | | 23_69 | 51_23 |
| Abau-H3-7 | 2006 | 1 | *baumannii* | Illumina | 3993785 | 66 | 82,8 | | 23_69 | 51_23 |
| Abau-H3-8 | 2006 | 1 | *baumannii* | Illumina | 3923677 | 80 | 44,3 | | 23_69 | 51_23 |
| Abau-H3-9 | 2006 | 1 | *baumannii* | Illumina | 3991877 | 65 | 45,5 | | 23_69 | 51_23 |
| Abau-H3-10 | 2006 | 3 | *baumannii* | Illumina | 3876967 | 39 | 44,2 | | 71 | 51 |
| Abau-H3-11 | 2006 | 3 | *baumannii* | Illumina | 3871471 | 39 | 70,0 | | 71 | 51 |
| Abau-H3-12 | 2006 | 25 | *baumannii* | Illumina | 4217980 | 45 | 44,6 | | 64 | 51 |

| Name | Year | STp | Species | Assembly type | Genome length | Contigs | Illumina coverage | ONT coverage | OXAs curated | OXA types curated |
|---|---|---|---|---|---|---|---|---|---|---|
| Abau-H3-13 | 2006 | 2 | *baumannii* | Illumina | 3978466 | 106 | 34,1 | | 66_72 | 51_24 |
| Abau-H3-14 | 2006 | 2 | *baumannii* | Illumina | 3823317 | 96 | 28,7 | | 66_72 | 51_24 |
| Agyl-H1-1 | 2004 | - | *gyllenbergii* | Illumina | 4731609 | 109 | 36,0 | | 671 | 286 |
| Ajoh-H1-1 | 2001 | - | *johnsonii* | Illumina | 3579359 | 133 | 42,5 | | 212 | 211 |
| Ajun-H1-1 | 2003 | - | *junii* | Illumina | 3670071 | 69 | 93,1 | | None | None |
| Ajun-H1-2 | 2004 | - | *junii* | Hybrid | 4025181 | 6 | 38,0 | 81,1 | 58 | 58 |
| Ajun-H1-3 | 2004 | - | *junii* | Hybrid | 3759138 | 6 | 79,7 | 22,7 | 58 | 58 |
| Alwo-H1-1 | 2003 | - | *lwoffii* | Illumina | 3199815 | 90 | 48,5 | | 1112 | 134 |
| Alwo-H1-2 | 2004 | - | *lwoffii* | Illumina | 3214352 | 195 | 68,2 | | 1110 | 134 |
| Alwo-H1-3 | 2004 | - | *lwoffii* | Illumina | 3504035 | 217 | 128,8 | | 1111 | 134 |
| Alwo-H1-4 | 2005 | - | *lwoffii* | Illumina | 3444482 | 122 | 115,0 | | 1112 | 134 |
| Apit-H1-1 | 2001 | - | *pittii* | Illumina | 3924444 | 84 | 43,5 | | 500 | 213 |
| Apit-H1-2 | 2004 | - | *pittii* | Illumina | 3832859 | 22 | 53,0 | | 1113 | 213 |
| Apit-H1-3 | 2004 | - | *pittii* | Illumina | 3916422 | 22 | 46,9 | | 1114 | 213 |
| Apit-H1-4 | 2004 | - | *pittii* | Illumina | 3918970 | 62 | 139,5 | | 506 | 213 |
| Apit-H1-5 | 2005 | - | *pittii* | Illumina | 3819466 | 19 | 55,2 | | 1115 | 213 |
| Apit-H1-6 | 2006 | - | *pittii* | Illumina | 3868698 | 115 | 40,6 | | 500 | 213 |
| Asch-H1-1 | 2003 | - | *schindleri* | Illumina | 3404196 | 121 | 59,1 | | 1116 | 134 |
| Aurs-H1-1 | 2001 | - | *ursingii* | Illumina | 3493960 | 93 | 57,1 | | None | None |
| Aurs-H1-2 | 2004 | - | *ursingii* | Illumina | 3441308 | 64 | 57,9 | | None | None |
| Avar-H1-1 | 2004 | - | *variabilis* | Illumina | 3427529 | 159 | 64,1 | | None | None |
